# A graph clustering algorithm for detection and genotyping of structural variants from long reads

**DOI:** 10.1101/2022.11.04.515241

**Authors:** Nicolás Gaitán, Jorge Duitama

## Abstract

Structural variants (SV) are polymorphisms defined by their length (>50 bp). The usual types of SVs are deletions, insertions, translocations, inversions, and copy number variants. SV detection and genotyping is fundamental given the role of SVs in phenomena such as phenotypic variation and evolutionary events. Thus, methods to identify SVs using long read sequencing data have been recently developed. We present an accurate and efficient algorithm to predict SVs from long-read sequencing data. The algorithm starts collecting evidence (Signatures) of SVs from read alignments. Then, signatures are clustered based on a Euclidean graph with coordinates calculated from lengths and genomic positions. Clustering is performed by the DBSCAN algorithm, which provides the advantage of delimiting clusters with high resolution. Clusters are transformed into SVs and a Bayesian model allows to precisely genotype SVs based on their supporting evidence. This algorithm is integrated in the single sample variants detector of the Next Generation Sequencing Experience Platform (NGSEP), which facilitates the integration with other functionalities for genomics analysis. For benchmarking, our algorithm is compared against different tools using VISOR for simulation and the GIAB SV dataset for real data. For indel calls in a 20x depth Nanopore simulated dataset, the DBSCAN algorithm performed better, achieving an F-score of 98%, compared to 97.8 for Dysgu, 97.8 for SVIM, 97.7 for CuteSV, and 96.8 for Sniffles. We believe that this work makes a significant contribution to the development of bioinformatic strategies to maximize the use of long read sequencing technologies.

## INTRODUCTION

Structural variants (SV) are a type of genetic polymorphisms, in both coding and non-coding sequences, which are usually defined by their length (>50 bp). The main types of SVs are deletions, insertions, translocations, inversions, and copy number variants (Alkan et al., 2011). The main genomic processes that cause the formation of structural variants are DNA recombination, replication, and repair-associated processes (Carvalho et al., 2016). For example, one common mechanism is Non-Allelic Homologous Recombination (NAHR) which is a genetic repair mechanism in which misalignment of previously duplicated regions called low copy repeats (LCR) occurs during meiosis. This subsequently causes a genomic rearrangement event on another locus that does not belong to the LCR gene, thus creating further deletions or duplications (Parks et al., 2015).

The interest in SVs comes mainly from the functional consequences of their genetic diversity. It has been proven that many SVs are involved in different gene expression patterns and influence different characteristics. SVs that are located adjacent to genes may affect *cis*-regulatory regions leading to either silencing or increasing gene expression, which can even explain a significant fraction of variation in Quantitative Trait Loci (QTL) (Chiang et al., 2017). Another case is when duplications increase the amount of overall transcript-protein production by gene dosage effect. This has proven to be beneficial for artificial selection in certain plant species where the average size of fruits increased because the plant variant suffered a specific duplication in a cytochrome coding gene (Alonge et al., 2020). A very interesting way in which SVs may affect expression is by causing structural changes, such as altering the position or composition of *cis*-regulatory regions. For example, Alonge *et al.* (2020) found that at least 50% of the SVs found in an assessment of around 100 lines of tomato were associated with gene expression regulatory processes, mostly reducing or silencing gene expression.

Structural variants also provide fundamental information about evolutionary relationships between organisms and their natural history. Many Whole-Genome Sequencing (WGS) studies have been conducted to assess the prevalence of different SVs and their variation in organisms, populations, or species. In plants, the analysis of structural variants allowed to elucidate the dynamics of whole-genome duplication (WGD) events and their evolutionary role (Qiao et al., 2019). WGDs are followed by a fast diploidization process, mainly because most of the duplicated genes become paralogs (Qiao et al., 2019). Furthermore, many components of the C4 metabolic pathway were brought by these WGD events and single duplication events. This is an interesting case of convergence throughout the evolution of different plant lineages (Wang et al., 2009). These changes are influenced by the synergistic effect of WGDs, transposed duplication, and dispersed gene duplication, evidenced by overlapping peaks in the rates of synonymous substitutions (Qiao et al., 2019). This shows how SVs can provide substantial amounts of evidence for evolutionary studies.

Given the importance of SVs, a large number of computational methods have been developed to identify and genotype SVs, based on high throughput sequencing (HTS) data. Most of these SV detection tools are based on short-read sequencing technologies (Cleal et.al., 2022; Sarwal et.al., 2022). This presents many limitations, mostly due to the length of structural variants, which usually exceeds the read length, which reduces the precision of both identification and genotyping (Luan et.al., 2020, Mahmoud et.al., 2019). Recently, new SV calling tools have adopted long reads as their input data, significantly increasing the accuracy of SV detection in comparison with short read based callers (Mahmoud et.al., 2019; Schwarz et.al., 2021). This has allowed many researchers to increase their catalog of functionally relevant structural variants, including some that affect the pathophysiology of diseases such as human cancer (Fujimoto et.al., 2021; Thibodeau et.al., 2020). However, further improvements could be achieved by novel algorithmic techniques. Some difficulties arise even when long reads are used. Since SV detection relies on accurate read alignment, dissimilar, partial or inaccurate read alignments obscure the signal to perform a consistent detection and genotyping of SVs. Thus, the results also depend on the accuracy of the aligner software (Heller et.al., 2019). Additionally, from a software design point of view, our experience indicates that most current tools are difficult to operate because they require a large number of specific libraries and versions, their implementations are not debugged correctly and exceptions are not handled appropriately. For short read based callers, these limitations have been described by a recent benchmark study by Sarwal et.al (2022).

Benchmarking of SV detection has been a difficult task to perform. There are few independently validated gold standard datasets for real sequencing data because experimental validation is difficult to perform at a large scale. Consequently, there is no consensus on which of the existing tools produces the closest result to a gold standard set. Bolognini et.al (2020) addressed this issue by implementing a simulation software called VISOR, which produces a complete haplotype-resolved sample genome and simulates read alignments from a list of SVs, with either Oxford Nanopore or PacBio error profiles. Trying to optimize the SV calling pipeline, Jiang et.al (2021) evaluated the accuracy of different SV callers using VISOR simulations on real reported human SVs. For the 20x simulated dataset, they report that the best tools are CuteSV (F1=0.8), SVIM (F1=0.798) and Sniffles (0.769). Additionally, they provide recommendations for SV calling best practices such as sequencing experiments with read lengths of about 20 kb at 20x depth. Regarding real datasets, the most widely recognised and best-curated case is the high confidence structural variant dataset (Sample HG002 on reference genome GRCh37) from the Genome In A Bottle human sample project (GIAB) crafted for benchmarking (Zook et.al.,2020). The events reported in this file come from a mixture of sequencing technologies and have been predicted by using a pipeline integrating many different tools.

Structural variant detection provides the possibility of finding biological insights with many different functional consequences. In this manuscript, we developed a new software solution that improves the detection of SVs from long read alignments using the DBSCAN algorithm to solve the clustering problem, and that implements a new bayesian genotyping model. This functionality is integrated into the bioinformatic software suite (NGSEP) to further facilitate the analysis of genomic data.

## RESULTS

### A new clustering algorithm for detection and genotyping of Structural Variants

Similar to previous algorithms, the process of structural variant detection and genotyping starts from reads aligned to a reference genome and is divided in three main stages described as follows.

#### 1. Signature *Collection*

The main input to this algorithm is a set of read alignments in SAM or BAM format, obtained from the alignment of long reads to an assembled reference genome. Signatures are individual signals of a structural variant that are contained within each read alignment. They can be divided into intra-alignment and inter-alignment signatures. Intra-alignment signatures consist of evidence of deletions or insertions that are predicted as part of the read alignment process. Thus, these signatures are collected reading the description of the alignment (encoded in the CIGAR field of the SAM format) to find signals of insertion or deletion. Conversely, reads with multiple discordant alignment segments, regarding their position or orientation, are selected to identify inter-alignment signatures. Inter-alignment signatures can provide evidence of all SV types. Signatures are filtered by the minimum length specified by the user and added to a collection, which is sorted by reference coordinates.

#### 2. Signature Clustering

Given a set of SV signatures, we implemented a graph-based clustering in which each cluster becomes a candidate SV event. A graph is built independently for each signature type. The vertices of the graph correspond to the input collection of signatures identified in the previous step. Each signature is represented by a tridimensional vector with three numeric values: Start coordinate in the reference genome (B_i_) end coordinate in the reference genome (E_i_), and signature length (L_i_). The cost m_ij_ of the edge between two signatures i and j corresponds to the Euclidean distance of their corresponding vectors:

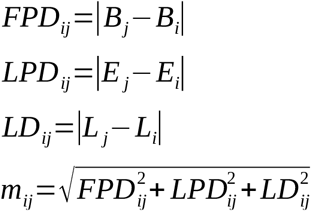

The DBSCAN algorithm is a non-supervised clustering procedure for n-dimensional vectors (points) based on the principle of density-based grouping (Schubert et.al., 2017). The parameters of this algorithm are a distance threshold (ε) to consider two points as density reachable (e.g. neighbors), and minimum number of neighbors (minPts) that a point should have to be considered a core point. Core points are fundamental to the process of delimiting the clusters, given that non-density reachable points from any core point are considered as noise or outliers. Thus, each cluster is defined by core points that are density reachable from each other. The algorithm traverses the points to identify their neighbors according to ε. Each point having at least *minPts* neighbors is declared as a core point. In this case, a new cluster is initialized with the core point and the neighbors as components and they are pushed to a queue where each node will also be queried for their neighbors, repeating this process accordingly. In other words, for each core point a Breadth First Search of the graph is performed until all of the density reachable points from any core point are visited, conforming a cluster. Figure 1 shows the main steps of this procedure.

**Figure 1.**
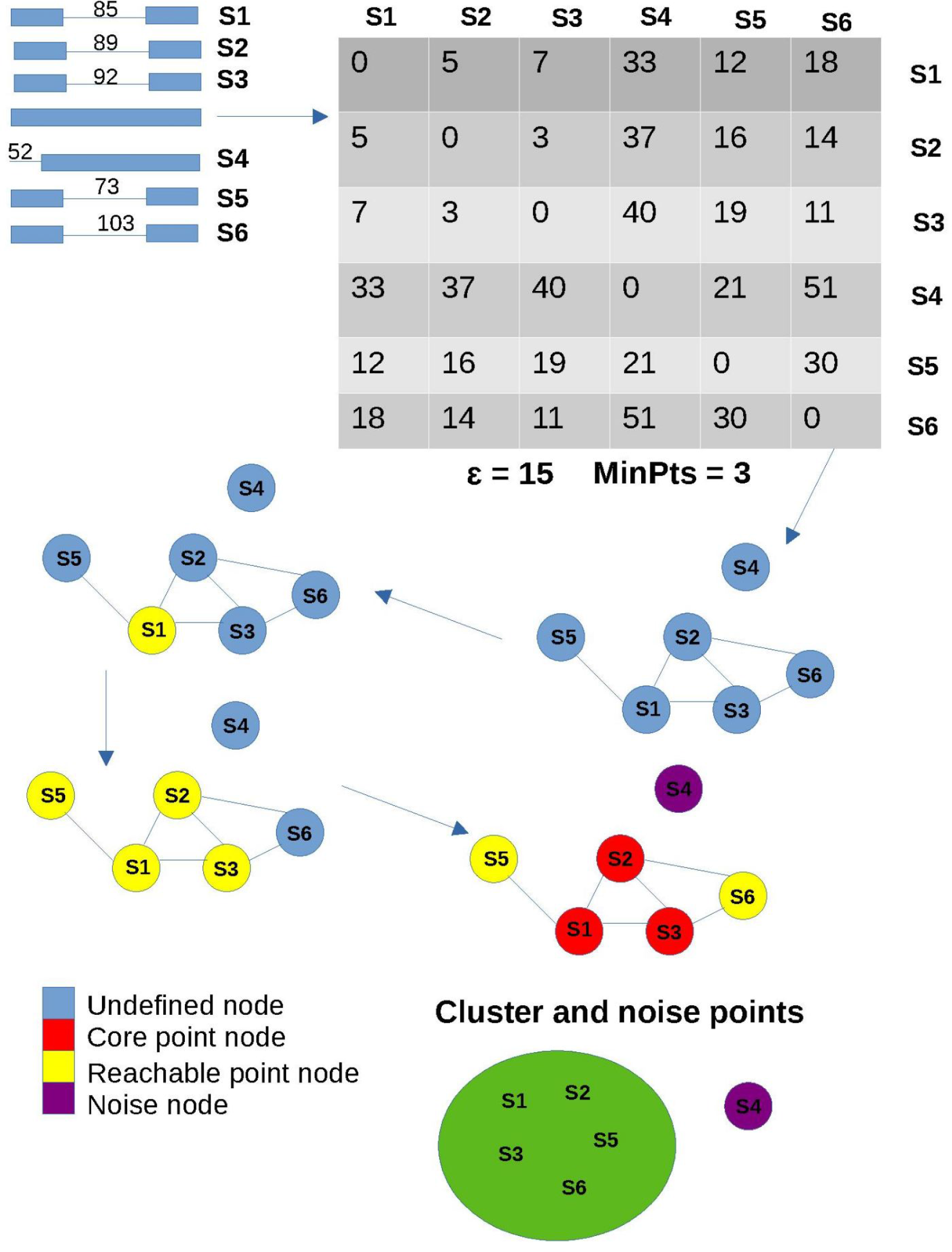
DBSCAN algorithm outlined in the context of variant calling. The process by which deletion signatures are clustered is shown. The noise point is not included in the cluster. The metric used in this case is euclidean distance based on three dimensions: The difference between the first and last reference position and the length of each signature.

#### 3. Cluster to Genotyped SV (SV-GT)

Each signature cluster identified in the previous step becomes a candidate SV. The last step of the process is the genotyping of these candidates. To identify the SV boundaries, the mean of the start reference coordinates of the signatures within the cluster is taken as the start coordinate of the SV. The end coordinate is calculated likewise. The length is taken as the difference between both averages, except for insertions where the mean of the cluster signatures is estimated. Candidate SVs are stored in a chromosome-position sorted collection. Then, a Bayesian genotyping process is performed for each candidate SV by reassessing the evidence that read alignments provide. To avoid having to reprocess the alignments file, a collection of simple alignments is kept in memory from the first stage, having the minimum possible information needed for this step. For each SV, intersecting read alignments are collected. Alignments supporting clustered signatures are considered evidence to support the alternative allele. If the spanning read alignment does not contain any evidence, it is counted as a call supporting the reference allele. Figure 2 shows the calculation of likelihood for the four scenarios generated by the combination of the two plausible alleles from which the read could be sequenced (hypotheses), and the two possible alleles that a read could support. The distribution of lengths of the signatures clustered to call the candidate SVs is used to calculate the likelihood of a read alignment supporting the SV. In this case, it is assumed that the read was actually sequenced from a chromosome affected by the SV (case 1). If a reference allele is assumed (case 2), a read with an SV signature is proposed to have happened by a misalignment error and a fixed value (0.0001 by default) is used as likelihood. The likelihood of a read supporting the reference allele that is assumed to be sequenced from a haplotype affected by the SV is calculated as the probability of having an indel error that reverts the SV and it is modeled by a gamma distribution (case 3). Finally, a fixed value (default 0.9999) is used for the likelihood of a read supporting the reference allele assuming sequencing from a reference haplotype.

**Figure 2.**
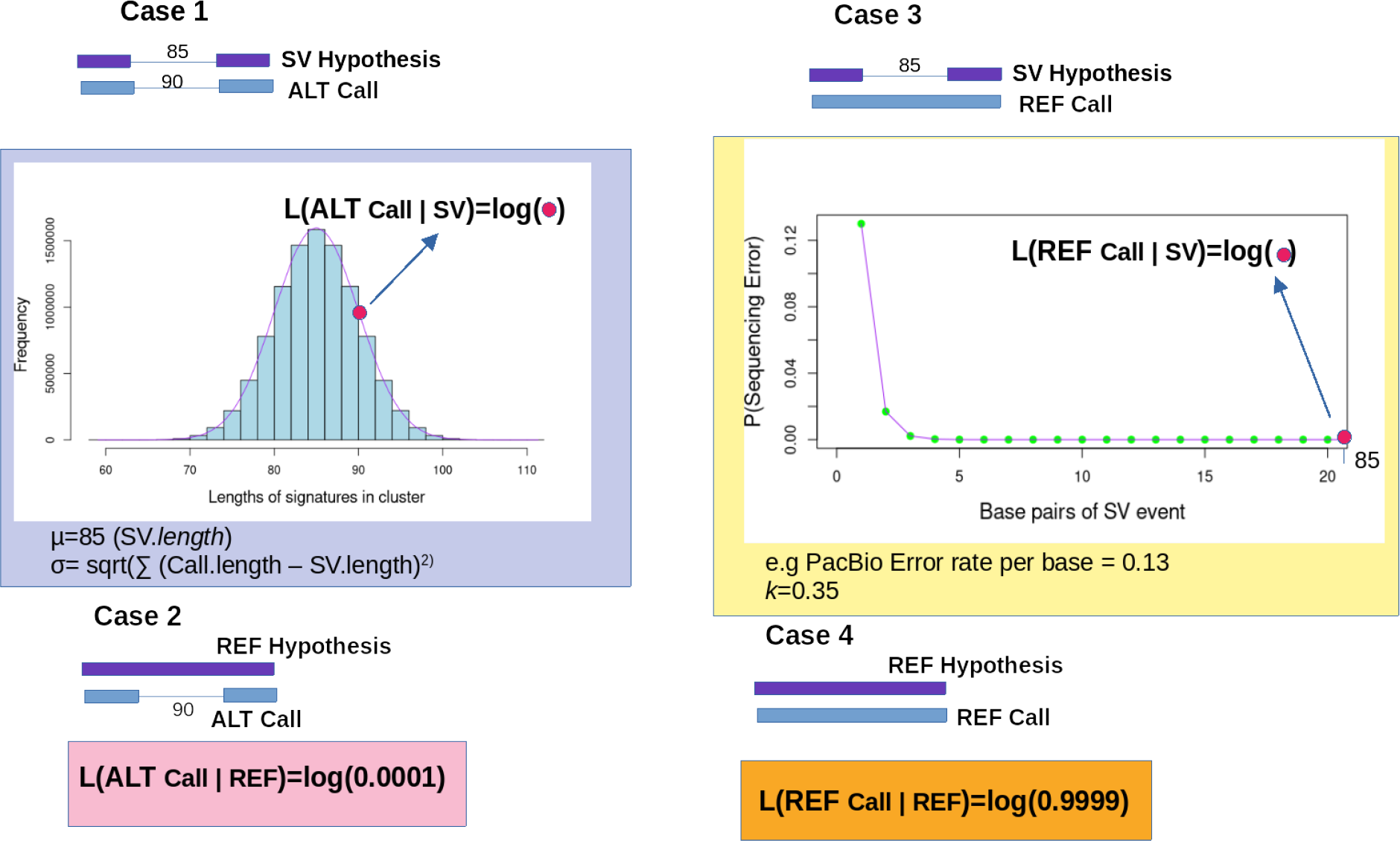
Likelihood estimation given each of four possible scenarios for a diploid organism. In each case, the base 10 logarithm of the obtained value is calculated.

Read likelihoods for each allele hypothesis are transformed into posterior probabilities for each possible genotype following the same procedure implemented to perform SNP genotyping (Gil et al., 2021). The hypothesis having the largest posterior probability is assigned as the predicted genotype. Similar to SNP genotyping, the quality of such SV calls will be the phred score *Q* corresponding to the genotype posterior probability.

### Benchmarking with simulation experiments

We performed two simulations of structural variants in the genome of *Arabidopsis thaliana* using the tool VISOR. This process generated 1718 insertions, 2532 deletions for the indel benchmark and 2065 inversions for the other one. The precision-recall results of our DBSCAN algorithm were compared to those of other tools, including SVIM (Heller et.al., 2019), Sniffles (Sedlazeck et.al., 2018), CuteSV (Jiang et.al., 2020) and Dysgu (Cleal et.al., 2022). After obtaining the metrics for both simulation experiments, precision-recall curves and F-score against depth were plotted for each tool. Additionally, execution times for each depth dataset were evaluated for single thread runs.

Figure 3A shows that the DBSCAN algorithm presented above outperforms all of the tools for depths of 20x,30x and 45x, producing calls with higher F-score. Only in the 60x test CuteSV outperforms DBSCAN by a small margin. Dysgu suffers from a significant decrease in precision, which causes the loss of F-score for bigger depths. DBSCAN shows a stable accurate performance regardless of alignment depth. The curve in figure 3B shows that while CuteSV and SVIM provide the best precision, DBSCAN maintains an almost equal value, while achieving a better recall for all tests. Meanwhile, Dysgu outperforms all of the tools in terms of recall for each coverage, but trades in precision, significantly dropping as depth increases. For inversions, SVIM produces the highest F-score, followed by the DBSCAN algorithm (Figure 3C).

**Figure 3.**
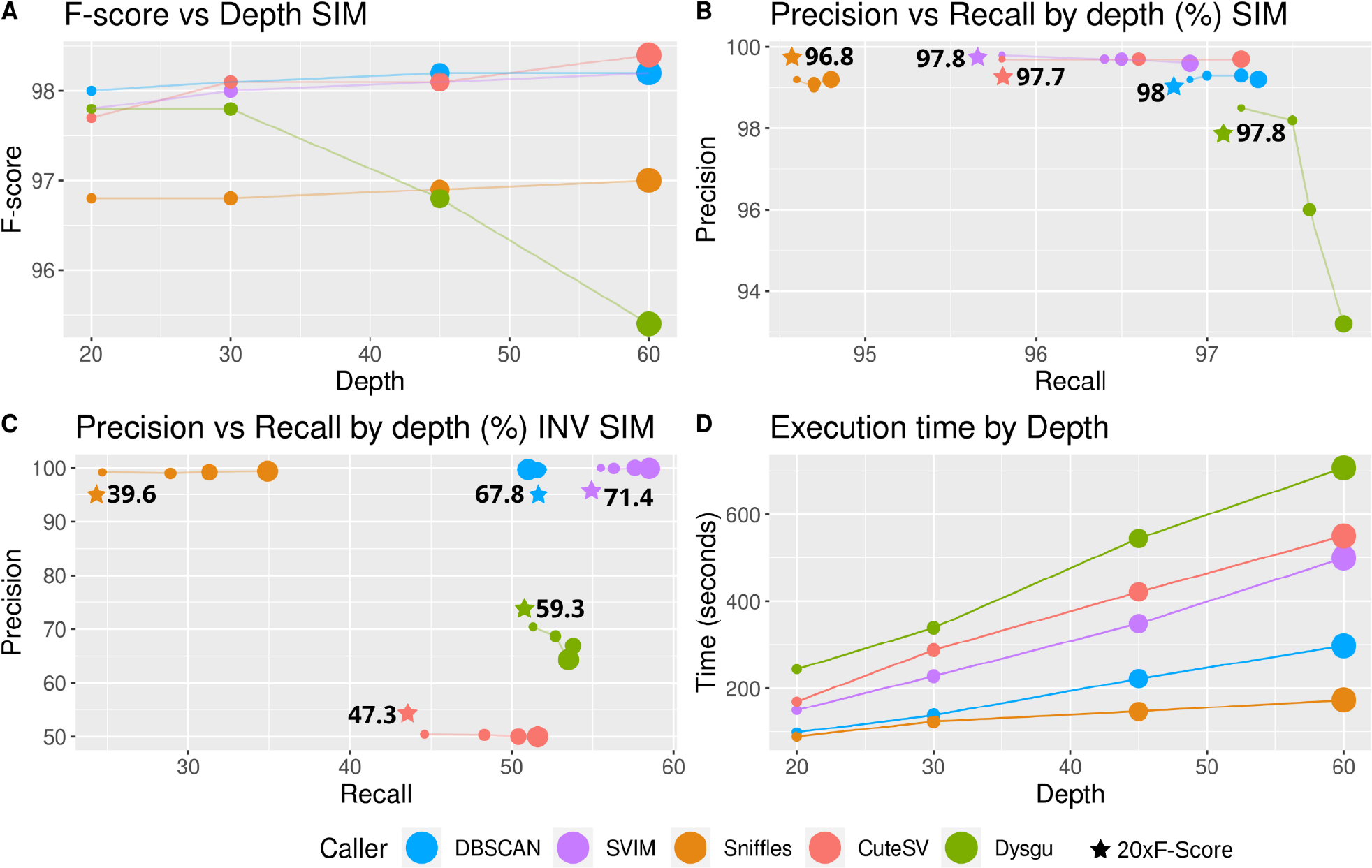
Simulation benchmarking results. The size of points increases according to depth for fixed values of 20x,30x,45x and 60x. A. F-score against Depth for all of the tools. For SVIM, a QS filter > 10 was applied given that this provides the best results for the tool, where 0 filter provides very low precision and >20 filters provide low recall. B. Precision-recall curve assessing values for varying depths of the input alignments. F-score values are shown for the 20x depth dataset for each tool. Again, SVIM datasets were filtered by 10 QS. C. The figure shows the same metrics as B but for the data simulated for inversions. D. Single thread execution time of all callers based on depth of the input alignments.

Single thread execution time was recorded for all experiments to compare the tools in terms of computational efficiency. As shown in figure 4D, all of them follow a linearly increasing trend. Sniffles and DBSCAN consistently required lower execution times compared to the other tools. It is worth clarifying that Sniffles, CuteSV and Dysgu support multithreading to reduce their execution times. Dysgu was the worst performing tool in terms of computational efficiency, running about three times slower than the DBSCAN algorithm.

**Figure 4.**
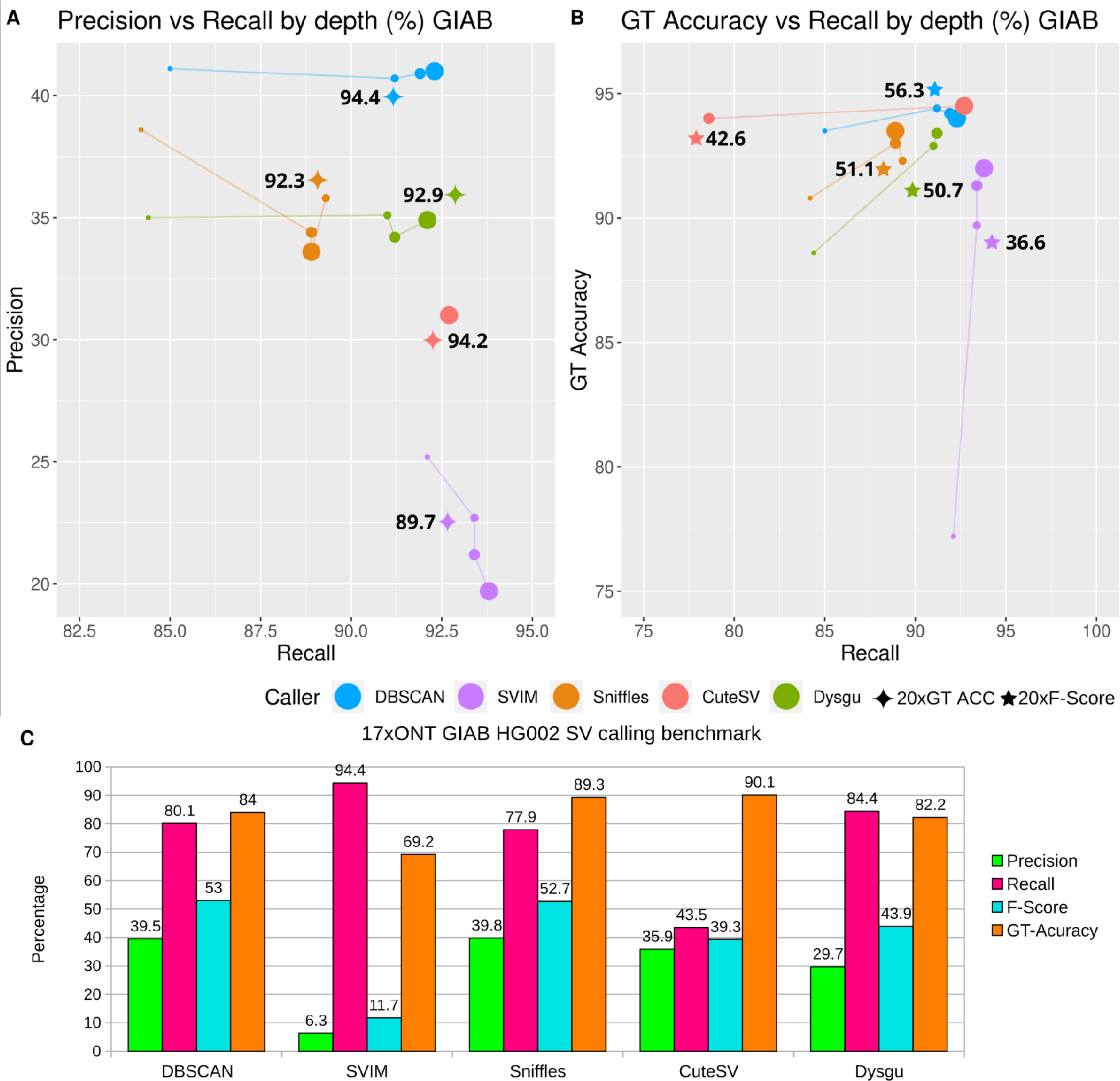
Performance metrics for PacBio HiFi and ONT data of varying depths from the HG002 sample variants with length > 50bp . For A,B the size of points increases according to depth for fixed values of 10x,20x,30x and 53x. A. Precision-recall curve by different depths. Genotyping accuracy is annotated for the 20x datapoint. B Genotyping accuracy against recall. The F-score is annotated at the 20x depth. C. Results of all benchmarking metrics on the 17x depth Oxford Nanopore reads dataset. For all results, GT Accuracy is calculated only for reported true positives.

### Benchmarking with the Genome In a Bottle human genome

Multiple experiments to determine recall, precision and genotyping accuracy of true positives were performed using the high confidence GIAB HG002 TIER 1 variants for chromosome 21 and 22 as the gold standard calls. First, the 53x PacBio CCS read collection coming from the HG002 subject was randomly sampled to 10x, 20x and 30x to perform varying depth benchmarking. Additionally, variant calling was performed on a 17x HG002 ONT. All metrics were obtained from the analysis performed with Truvari (English et.al., 2022).

For the PacBio HiFi varying depth experiments, Figure 4A shows that the best performing tool in terms of precision is DBSCAN. It is important to state that this metric is underrepresented given that the gold standard dataset only includes the most reliable calls in the HG002 dataset (Filter = PASS, length > 50). The recall of the DBSCAN algorithm is also superior to that of the other tools, except SVIM, which achieves the best recall using the entire dataset (53x) at the expense of a very low precision. Regarding genotyping accuracy, CuteSV calls achieve the highest value in this metric, closely followed by DBSCAN (Figure 4B). However, the DBSCAN algorithm is more robust to changes in the average read depth. Regarding the 17x ONT results, the same underrepresentation of precision happens, but it is consistent with the CCS results, where DBSCAN provides the best value and SVIM the worst in exchange for the best recall (Figure 4C). For GT accuracy, the best tools are Sniffles and CuteSV, but only the former provides good results in the other metrics. In general, the most consistent results for both types of HTS data and varying depths are provided by DBSCAN, shown by the highest F-score and a very good GT-accuracy.

## DISCUSSION

The availability of long read sequencing technologies represented a big step forward towards an accurate identification and genotyping of structural variants (Fujimoto et.al., 2021; Thibodeau et.al., 2020). Achieving this goal is becoming a requirement for current genomics, given the documented role of SVs as drivers of phenotyping variability and evolution (Alonge et al., 2020; Gorkovskiy et.al., 2021; Qiao et al., 2019; Wang et al., 2009). In this work we present the results of our efforts to develop novel algorithmic techniques, aiming to increase accuracy of both discovery and genotyping of SVs. Transforming the problem of clustering SV signatures into a geometric clustering problem in an Euclidean space, allowed us to integrate the well known DBSCAN clustering algorithm (Schubert et.al., 2017) to identify SVs. Integrating previous experiences implementing Bayesian models for SNV genotyping, allowed us to increase the accuracy of SV identification and provided a framework for SV genotyping. Benchmarking experiments running simulations and analyzing the GIAB dataset indicate that our algorithm achieves superior accuracy compared to current software solutions.

Researchers performing population genomic studies usually trade read depth by number of samples sequenced, looking for a balance that maximizes the cost-benefit of the sequencing effort (Cericola et.al., 2018; Fumagalli et.al., 2013). Thus, it is extremely important for SV detection tools to be able to produce accurate results from a low depth input. One of the biggest advantages of the DBSCAN algorithm when compared to the other state-of-the-art tools is that it is robust to reductions of read depth. Even at 20x average read depth, the integration of the Bayesian model provided the best results for genotyping accuracy. This outcome was also observed in the analysis of the 17x ONT reads, which have a bigger error rate than CCS reads. This suggests that our algorithm is also robust to increased per-base error rates. Beyond tools comparison, our experiments indicate that an average read depth around 20x is sufficient to achieve high detection and genotyping accuracy.

We believe that this work represents a significant contribution to current research on algorithms to analyze long DNA sequencing reads. We expect that the new functionality developed in NGSEP for SV detection from long reads will be useful for a large number of ongoing and upcoming research in population genomics for different species.

## METHODS

### Software development and integration within NGSEP

The algorithm described in this manuscript was implemented in Java 11 as a new option of the single sample variants detector functionality of the NGSEP software tool. Reuse of different NGSEP classes significantly decreased the development effort needed to code. Initially, for computing the input file a ReadAlignment iterator found in the ReadAlignmentFileReader class was used, given that it already collects all of the necessary information for each alignment. A Collection interface class named GenomicRegionSortedCollection allowed GenomicVariant interface implementing objects, such as Signature and CalledGenomicVariant objects, to be stored by sorted sequence, e.g chromosomes, and by genomic position. This also facilitated computed spanning alignments to specific variants. Additionally, the work made for genotyping SVs consisted mostly of programming the functionality to estimate likelihoods, given that the class CountsHelper allowed to calculate the genotype posterior probabilities, as it was implemented before to genotype small variants and SNPs. The class diagram for the functionality inside of the NGSEP class context is shown in figure 5.

**Figure 5.**
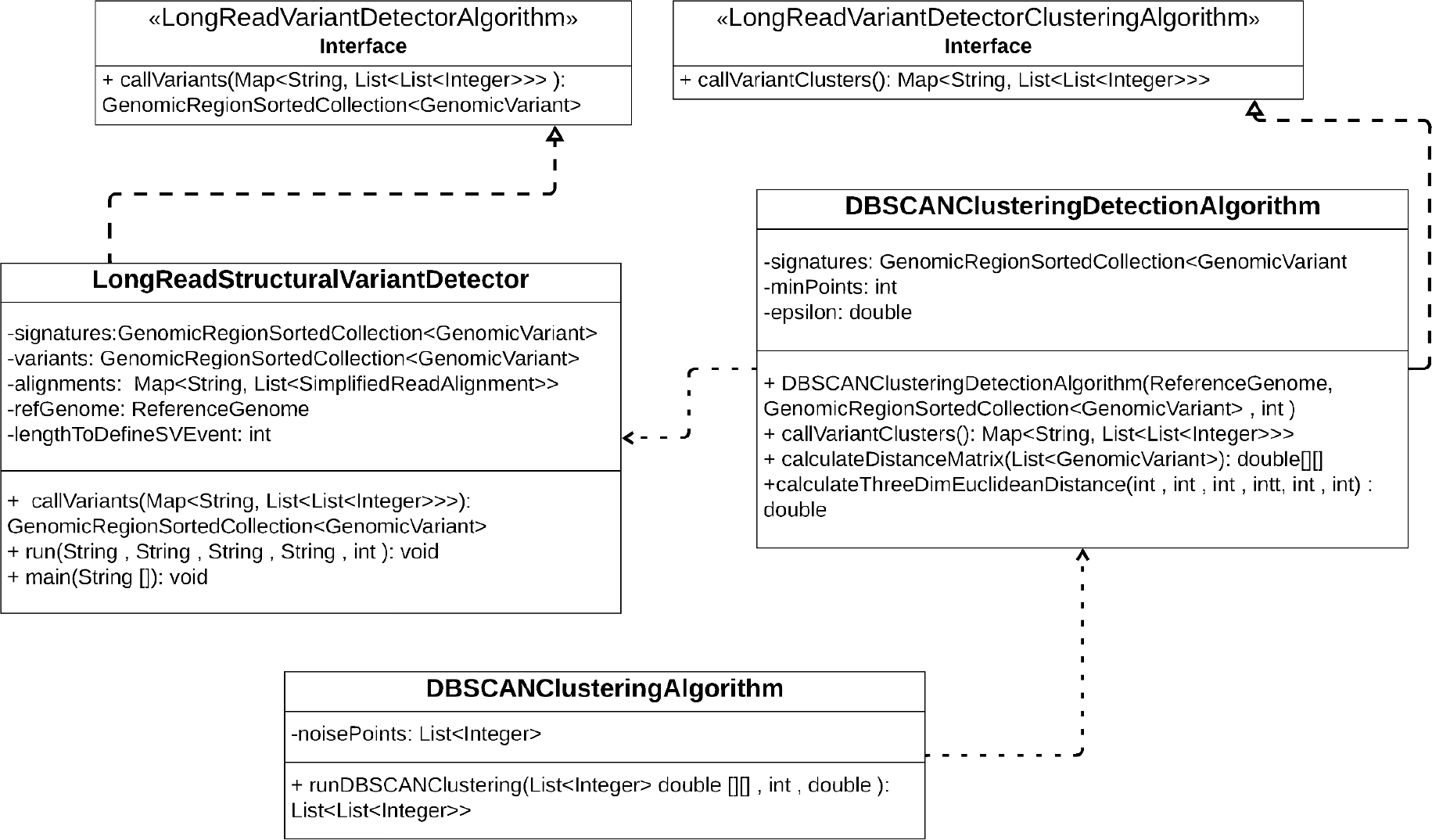
Class diagram of the functionality as a UML integrated in NGSEP. The generic algorithm classes are designed so that they can be used for other problems and applications. The clustering detection classes adapt them for the SV clustering functionality dictated by an interface designed to guide the solution to this problem.

### Simulation experiments

In order to assess the improvements this functionality presents in the SV calling problem, a thorough benchmarking process was established to evaluate performance metrics of recall, precision and efficiency. After an in-depth literature revision, four tools were included in the benchmark based on their performance and impact, including SVIM (version 2.0.0) (Heller et.al., 2019), Sniffles 2 (version 2.0.6) (Sedlazeck et.al., 2018), CuteSV (version 1.0.13) (Jiang et.al., 2020) and Dysgu (version 1.3.11) (Cleal et.al., 2022). Two strategies were chosen to produce a robust comparison, between simulations and real data. To obtain the established performance metrics for both approaches, the output VCF files were compared to the Gold Standard files using the software Truvari (English et.al., 2022), which provides recall, precision, f-score and genotype accuracy of the true positive calls. This tool has been recommended for benchmarking of genomic variant callers (Zook et.al., 2020).

SVs were simulated with the software VISOR (Bolognini et.al., 2020), based on the *Arabidopsis thaliana TAIR10* reference genome (Lamesch et.al., 2012). A total of 4,330 indel structural variants with a minimum length of 50bp were simulated (2500 deletions, 1830 insertions), and a genome containing these variants was generated. Next, reads with the characteristics of the Oxford Nanopore Sequencing Technology (ONT), including the error profile, were simulated with VISOR from this altered genome. Reads were aligned to the original reference genome using minimap2 (Li et.al., 2018). This pipeline was repeated to simulate four datasets of varying depths, including 20x, 30x, 45x and 60x. The resulting alignments were used as the input data for all tools.

### GIAB high-confidence dataset

The Genome in a Bottle (GIAB) consortium has produced a high confidence curated SV dataset, consisting of around 12.000 structural variants called from many different sequencing technologies and discovered by different bioinformatic methods on the Ashkenazi son sample (HG002) compared to the GRCh37 reference genome (Zook et.al., 2020). To perform benchmarking based on this asset, indel variants of more than 50bp in length, which fulfill the PASS filter (the most reliable calls) and that belong to the chromosome 21 and 22 were extracted as the reference Gold Standard. All the aforementioned software were run on a PacBio CCS HiFi read dataset of around 53x depth of the same sample. The same alignments were subsetted to produce 10x, 20x and 30x input files, in order to assess the effect of depth variance on the calling algorithms. A 17x depth Oxford Nanopore alignment dataset was constructed from the same HG002 sample to evaluate the effect of the long read sequencing technology on the SV calling process, considering the error rate of ONT reads is higher than CCS reads. Truvari (English et.al., 2022) was again used to produce the benchmark metrics, using symbolic alleles only, similar to the simulated data benchmark.. Minimap2 (Li et.al., 2018) was used to align all reads to the reference genome.

## ACKNOWLEDGEMENTS AND FUNDING

This work has been supported by the “Patrimonio autónomo del Fondo Nacional de Financiamiento para la ciencia, la tecnología y la innovación Francisco José de Caldas” with the contract number 80740-441-2020, awarded by the Colombian Ministry of Science to JD. We also acknowledge the high performance computing unit of Universidad de Los Andes for their technical support to conduct the benchmark experiments presented in this manuscript.

## DATA AVAILABILITY

The algorithm presented in this study can be executed through the Single sample Variants Detector functionality of the open source software Next Generation Sequencing Experience Platform (NGSEP). Releases of NGSEP are available at sourceforge (http://ngsep.sf.net). Life development is available in github (https://github.com/NGSEP).

The *A.thaliana* TAIR10 reference genome used for simulations is available in the phytozome v.12 database (https://phytozome-next.jgi.doe.gov). The GIAB TIER1 SV gold standard VCF file can be downloaded from the GIAB web site (https://www.nist.gov/programs-projects/genome-bottle). The GHC37 human reference genome can be found at the NCBI Assembly database (accession number GCA_000001405.1). PacBio HiFi and Oxford nanopore raw reads were downloaded from the NCBI SRA database (accession ids **<u>SRX7083056</u>**, and **<u>SRX9363340</u>** respectively).

